# SIREN: Predicting RNA-Small Molecule Interaction using Language Models without Requiring Structure

**DOI:** 10.64898/2026.11.19.768003

**Authors:** Wonkyeong Jang, Yein Kim, Jinseo Heo, Yeongju Lee, Woong-Hee Shin

## Abstract

RNA-targeted drug discovery is constrained by the scarcity of resolved RNA tertiary structures and the conformational flexibility of RNA, which limit the applicability of structure-based virtual screening. Here, SIREN (Small molecule Interaction with RNA via sEquence-based predictioN) is presented, a sequence-based framework that ranks small molecules for a given RNA target directly from its primary sequence without requiring tertiary structure. SIREN integrates a pretrained RNA language model with molecular language model embeddings and physicochemical descriptors through an interaction network that outputs a target-specific compound-ranking score. On four held-out ROBIN riboswitch targets, SIREN outperforms the structure-based methods RNAmigos2, RNAmigos++, and rDock in both enrichment and overall discrimination, and a cross-target derangement analysis confirms that its rankings depend on RNA identity rather than general ligand propensity alone. On an independent HIV-1 TAR benchmark, SIREN achieves strong discrimination without an experimentally derived RNA conformational ensemble. Prospective screening against IRES-derived RNA hairpins from cMyc and XIAP mRNAs further identifies multiple experimentally validated, micromolar-affinity binders from a diverse chemical library. These results demonstrate that SIREN generalizes to unseen RNA targets, transfers across independent benchmarks, and provides a scalable, structure-free framework for RNA-targeted compound prioritization that complements existing structure-based screening approaches.

## 1. Introduction

RNA plays diverse roles in gene regulation, translation, and structural organization, and is implicated in various human diseases, including cancer and neurological disorders^[1–3]^. Because only a small fraction of the human genome encodes proteins, and the majority of those proteins are considered undruggable, there has been growing interest in developing drugs that target RNA^[3–5]^. Small molecules have been successfully developed against ribosomal RNAs, riboswitches, and structured viral RNA elements^[3,6,7]^. Risdiplam represents a clinically validated example of targeting human RNA directly with small molecules^[8]^. Despite this progress, computational modeling of RNA–small-molecule interactions remains challenging due to limited structural data and the conformational complexity of RNA^[9]^. Effective computational screening approaches are therefore needed to support RNA-targeted drug discovery.

Existing computational approaches for small-molecule drug discovery targeting RNAs span both traditional scoring methods and machine learning-based models. SPRank^[10]^ is a knowledge-based atomic pairwise scoring function that uses a random forest regressor to evaluate docking poses, incorporating stacking and solvation-related effects to score a given RNA-ligand complex. RLDOCKScore^[11]^ rescores docking poses by introducing explicit terms for base-ligand stacking interactions, ionic screening, and ligand conformational entropy. This improvement in virtual screening is achieved while retaining native-pose prediction capability. RNAmigos2^[12]^ is a program that utilizes 2.5D graphs to depict small-molecule binding sites. It employs a combination of self-supervision and synthetic data derived from docking to facilitate structure-based virtual screening with competitive enrichment. RNAsmol^[13]^ adopts a structure-free, sequence-based framework that integrates data perturbation and augmentation with multiview convolution, graph diffusion convolution, and attention-based fusion to enhance interaction classification, unseen evaluation, and decoy ranking. DeepRNA-DTI^[14]^ further extends sequence-based modeling through a multitask architecture that jointly predicts RNA–compound interactions and nucleotide-level binding sites using pretrained RNA and molecular representations. SMRTnet^[15]^ eliminates the necessity for RNA tertiary structures by integrating RNA sequence and secondary structure with small-molecule SMILES and graph-derived features through multimodal fusion. The program identified experimentally validated hits in prospective screening. Notwithstanding the advances achieved thus far, contemporary methodologies continue to entail compromises among three fundamental aspects: reliance on RNA tertiary structures, screening efficiency, and generalization across uncharted RNA targets and chemical space. In particular, structure-based methods are constrained by the availability and quality of RNA tertiary structures and by docking-related computational cost. On the contrary, sequence-based methods alleviate these constraints but still leave room for improvement in robust generalization and in the effective integration of complementary RNA and ligand representations^[16]^.

In order to address the aforementioned limitations, we present SIREN (Small molecule Interaction with RNA via sEquence-based predictioN), a sequence-based RNA-ligand virtual screening framework without requiring RNA tertiary structures. SIREN integrates pretrained RNA language model embeddings with molecular language model representations and descriptor-based ligand features, thereby enabling complementary characterization of RNA context and ligand properties within a unified predictive framework. SIREN has been developed to enable efficient and scalable RNA-targeted virtual screening. This objective is achieved by preserving rich multimodal molecular information while eliminating explicit structural dependence on RNA. Following its training on the ROBIN dataset^[17]^, the program demonstrated superior performance in screening RNA-targeting molecules on the benchmark test when compared to structure-based methods. SIREN was further benchmarked to the human immunodeficiency virus type-1 transactivation response element (HIV-1 TAR). In addition, the program was also applied to screen compounds against the two RNA hairpins derived from the internal ribosome entry sites (IRESs) in the 5′ untranslated regions of cMyc and XIAP mRNAs, and prioritized compounds were experimentally validated for binding to the corresponding RNA constructs. The findings demonstrate the capacity of SIREN to generalize to unseen RNA targets and transfer to an independent virtual screening benchmark. Furthermore, the study substantiates its utility through prospective screening and experimental validation of prioritized compounds.

## 2. Results

### 2.1. Overview of SIREN

SIREN is a sequence-based virtual screening framework designed to prioritize small molecules for a given RNA target without requiring tertiary structures (**Figure 1**). The framework represents RNA directly from its primary sequence using a pretrained RNA language model and represents each candidate ligand using a combination of molecular language model embeddings and physicochemical descriptors. The contextualized nucleotide-level RNA embeddings are summarized into a fixed-dimensional target representation through attention-based pooling, while the ligand features are projected into a corresponding latent representation. The RNA and ligand representations were then integrated by an interaction network that captures both molecule-specific features and RNA-ligand compatibility. The resulting projection is transformed into an interaction score ranging from zero to one, which is then used to rank compounds for each RNA target. Higher scores indicate a greater predicted propensity for interaction.

**Figure 1.**
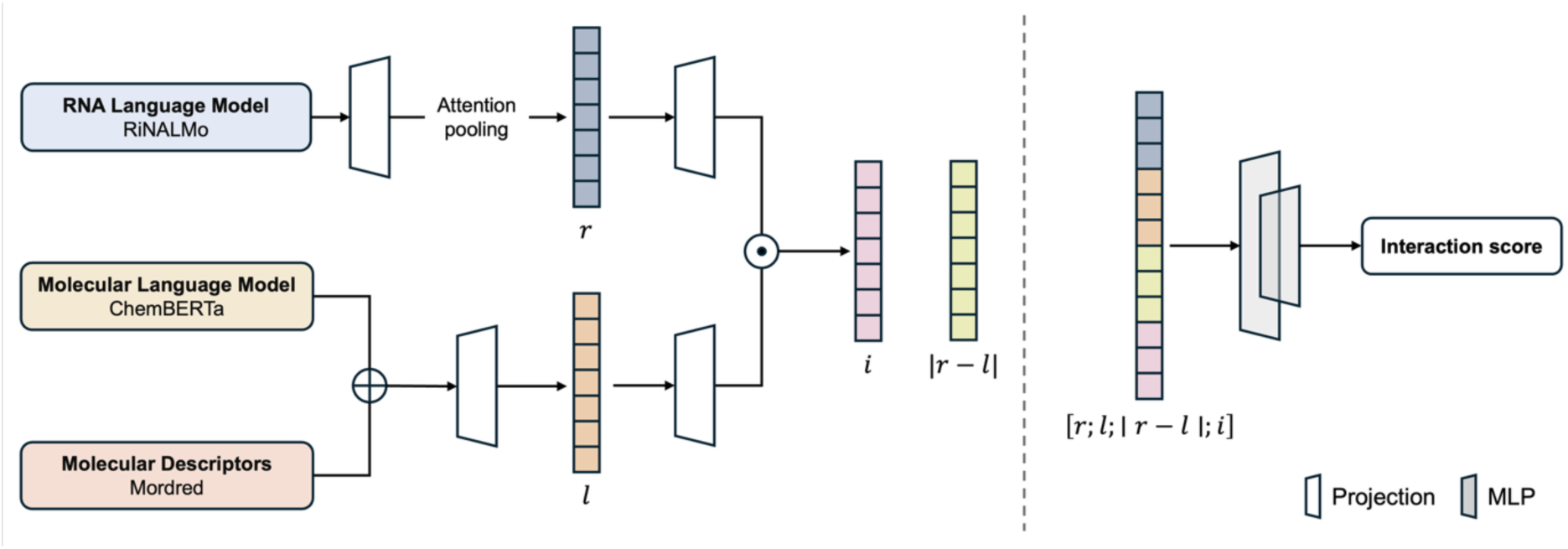
Overview of the SIREN architecture. RNA sequences are encoded using a pretrained language model and summarized by attention-based pooling, while ligands are represented using molecular language model embeddings and molecular descriptors. The resulting RNA and ligand representations are projected into a common latent space and integrated through an interaction network to generate a target RNA-specific compound-ranking score.

The RNA and ligand representations for SIREN were systematically investigated. A set of three pretrained RNA language models—RiNALMo^[18]^, ERNIE-RNA^[19]^, and RNA-FM^[20]^—was evaluated in combination with eight ligand representation schemes. These schemes were constructed from two compound language models, ChemBERTa^[21]^ and MolFormer^[22]^, and two descriptor sets, RDKit^[23]^ and Mordred^[24]^. This process yielded 24 RNA-ligand representation configurations, which were evaluated using a consistent prediction architecture, data partitioning strategy, and training procedure on the RNA subset of the ROBIN dataset small-molecule microarray (SMM) dataset^[17]^. To reduce sequence redundancy between the training and validation subsets, RNA targets were partitioned at the RNA-cluster level. Given that active compounds constituted a minority of the screening data, the primary criterion for model selection was validation area under the precision-recall curve (AUPRC). The RiNALMo-ChemBERTa with Mordred configuration obtained the highest validation AUPRC and was consequently selected as the ultimate SIREN model for ensuing analyses.

Subsequently, we evaluated the capacity of the selected model to generalize beyond the RNA targets utilized during its development. Four ROBIN riboswitch targets with available crystal structures—PreQ1, SAM-II, TPP, and ZTP—were excluded from training and validation and retained as a fixed held-out benchmark. These targets were utilized to evaluate the discrimination and early enrichment of unseen RNA sequences, and to compare SIREN with previously reported virtual screening methods RNAmigos2, RNAmigos++^[12]^, and rDock^[25]^. To determine whether the predicted rankings reflected RNA-specific information rather than general preference for particular ligands, we additionally compared the correctly matched RNA-ligand score assignments with complete cross-target derangements using compounds shared across the four targets.

To assess the transferability of SIREN beyond the ROBIN assay setting, the study further evaluated SIREN on an independent HIV-1 TAR benchmark. This benchmark contained experimentally characterized binders and non-binders^[26]^. The performance of this dataset was compared with previously reported docking and scoring approaches, including methods based on experimentally informed TAR conformational ensembles. Finally, the prospective screening utility of SIREN was assessed by ranking 2,333 compounds from ChemBank of Korea Research Institute of Chemical Technology (KRICT) against RNA hairpins derived from the 5′ UTR IRESs of cMyc and XIAP mRNAs. For each RNA target, the top-ranked 233 compounds, corresponding to 10% of the library molecules, were selected for experimental validation.

### 2.2 Evaluation and Selection of RNA and Ligand Configuration for SIREN

To determine which representations most effectively capture RNA target and ligand information for virtual screening, we systematically evaluated 24 combinations of three RNA language models and eight ligand representation schemes. The objective of this analysis was twofold: first, to assess the contribution of the RNA encoder, and second, to determine whether pretrained molecular embeddings and physicochemical descriptors provide complementary ligand information. Given that active compounds constituted a minor proportion of the screening dataset, AUPRC, which emphasizes retrieval of the positive class, was utilized as the primary metric for model selection, with the area under the receiver operating characteristic curve (AUROC) reported as a complementary measure. The results of the study are summarized and illustrated in **Figure 2**. The detailed results are given in **Table S1**.

**Figure 2.**
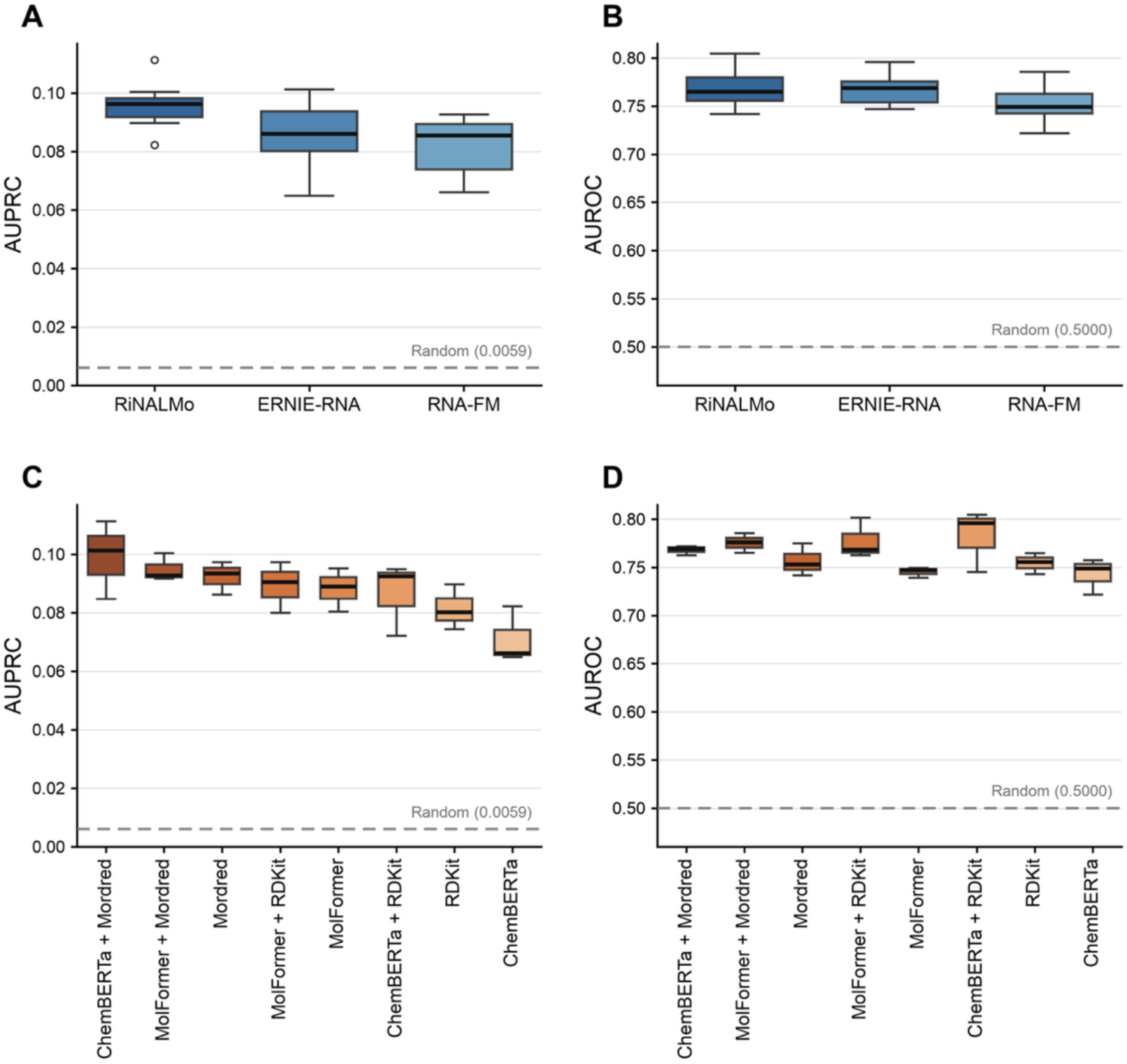
Validation Performance across all RNA and ligand representations. (A, B) AUPRC and AUROC grouped by RNA language model; (C, D) corresponding values grouped by ligand representations. Box plots show the median, interquartile range, and 1.5 times the interquartile range across alternative representation combinations. Categories are ordered by median validation AUPRC. The gray dashed lines indicate random-selection baselines.

For reference, the expected AUPRC under random compound ranking, averaged across the 23 RNA targets used for model training and validation, was 0.0059. The RNA representation exerted a substantial influence on the predictive performance across ligand representations. It was demonstrated that RiNALMo achieved the highest mean validation AUPRC across the eight ligand configurations (0.096), followed by ERNIE-RNA (0.086) and RNA-FM (0.082) (Figure 2A). RiNALMo also demonstrated the highest mean AUROC (0.770), although the discrepancy from ERNIE-RNA (0.767) was negligible. RNA-FM yielded a lower mean AUROC of 0.752 (Figure 2B).

The impact of ligand representation on performance was also significant. The highest mean AUPRC across the three RNA language models was achieved by ChemBERTa+Mordred (0.099), followed by MolFormer+Mordred (0.095) and Mordred alone (0.092) (Figure 2C). In contrast, the highest mean AUROC was obtained with ChemBERTa+RDKit (0.782), followed by MolFormer+RDKit (0.778) and MolFormer+Mordred (0.776) (Figure 2D). The efficacy of molecular language model embeddings in terms of AUPRC was found to be inferior, with ChemBERTa exhibiting the lowest mean AUPRC of 0.071. Overall, descriptor-containing representations tended to achieve higher AUPRC than molecular language model embeddings alone. This finding suggests the presence of information that is not fully captured by the pretrained molecular representations, and that is potentially provided by explicit physicochemical features.

Among the 24 individual configurations, RiNALMo combined with ChemBERTa+Mordred achieved the highest validation AUPRC (0.111), with an AUROC of 0.772. The combination of RiNALMo with ChemBERTa+RDKit yielded AUROC of 0.805, yet its AUPRC was lower at 0.092. Given the strong class imbalance and the virtual screening objective of enriching active molecules among top-ranked candidates, the former was selected as the final SIREN configuration. This selected configuration therefore represents an approximately 19-fold enrichment over the random-selection baseline AUPRC established above (0.0059).

### 2.3 Evaluation of SIREN on Held-out RNA Targets

To assess the model’s capacity for generalization to unseen RNA targets, the selected RiNALMo-ChemBERTa+Mordred model was evaluated on four ROBIN riboswitch targets—PreQ1, SAM-II, TPP, and ZTP—that were completely excluded from model development. For each target, compounds were then assigned a ranking based on the interaction scores predicted by SIREN. The performance of the method was evaluated using AUROC and enrichment factors (EFs) at the top 1%, 2%, and 5% of each ranked compound library. These results were then compared with the reported benchmark results for RNAmigos2, RNAmigos++, and rDock, structure-based virtual screening methods (**Table 1** and **Table S2**).

**Table 1.**
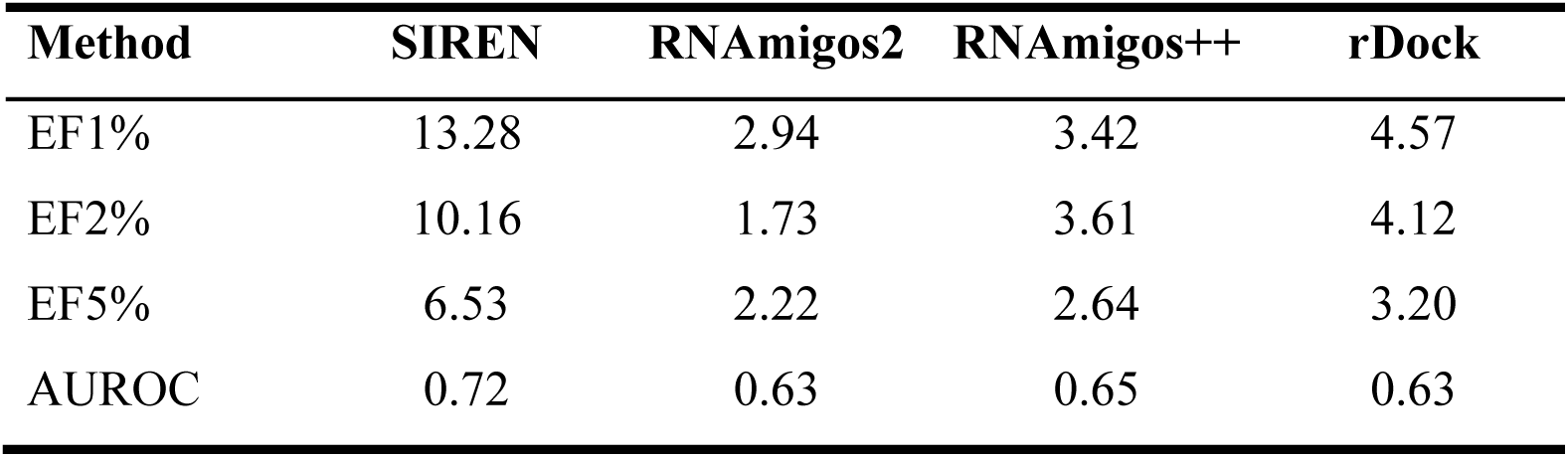
Virtual-screening performance on four held-out ROBIN RNA targets. Enrichment factors at the top 1%, 2%, and 5% of the ranked compound library and AUROC are reported across PreQ1, SAM-II, TPP, and ZTP. Comparator values are taken from the RNAmigos2 study^[12]^.

| **Method** | **SIREN** | **RNAmigos2** | **RNAmigos++** | **rDock** |
|---|---|---|---|---|
| EF1% | 13.28 | 2.94 | 3.42 | 4.57 |
| EF2% | 10.16 | 1.73 | 3.61 | 4.12 |
| EF5% | 6.53 | 2.22 | 2.64 | 3.20 |
| AUROC | 0.72 | 0.63 | 0.65 | 0.63 |

Across the four held-out targets, SIREN achieved a mean AUROC of 0.72, compared with reported values of 0.65 for RNAmigos++, and 0.63 for RNAmigos2 and rDock. The mean EF1%, EF2%, and EF5% values for SIREN were 13.28, 10.16, and 6.53, respectively. The highest reported comparator (rDock) values at the same cutoffs were 4.57, 4.12, and 3.20. These values correspond to 2.91-, 2.47-, and 2.04-fold increases over the best comparator at each cutoff. It is noteworthy that the advantage was most evident at the earliest screening cutoffs, where enrichment is especially pertinent because only a limited number of top-ranked compounds can typically be advanced to experimental testing.

The performance exhibited consistency across all four held-out targets, rather than being influenced by a specific target. The AUROC values ranged from 0.67 to 0.77, while the EF1% ranged from 9.14 to 16.02. Similarly, the EF2% varied from 8.22 to 12.33, and EF5% demonstrated a range of 5.24 to 8.23. SIREN demonstrated the highest AUROC for TPP at 0.77, whereas the highest EF1% was observed for ZTP (16.02). Together, these results suggest that the selected model for SIREN maintained its discriminatory and early-enrichment capabilities across RNA sequences that were not encountered during the model development.

Subsequently, we investigated whether ligand rankings depended on RNA sequence information, or whether they merely reflected a target-independent propensity of particular compounds to bind RNA. This analysis encompassed 14,302 ligands shared across the four ROBIN targets, corresponding to 57,208 RNA–ligand pairs and 485 positive interactions. The RNA–score assignments that were correctly matched were then compared with all nine complete cross-target derangements, in which no RNA retained scores generated for itself. Utilizing within-target percentile ranks, the correctly matched assignment attained a pooled AUROC of 0.75, in comparison to a mean of 0.72 across the derangements, corresponding to a discrepancy of 0.024 (bootstrap 95% CI, 0.010–0.039; one-sided *P*=0.001). Similarly, the EF1% exhibited a decline from 14.41 for the accurate assignments to an average of 12.97 following cross-target reassignment, making a discrepancy of 1.441 (bootstrap 95% CI, 0.284–3.078; one-sided *P*=0.011). The reduction in both AUROC and early enrichment after RNA-score assignment indicates that SIREN rankings depend in part on the identity of the RNA target rather than being explained solely by general ligand propensity. The moderate magnitude of this effect further suggests that the predictions contain both RNA-dependent information and a target-independent component associated with ligand propensity.

### 2.4 Robust Performance of SIREN on an Independent HIV-1 TAR benchmark

The screening performance of SIREN was further validated using an independent virtual screening for the HIV-1 TAR. HIV-1 TAR is a structured RNA element formed at the 5’ end of nascent viral transcripts, where it binds the viral transactivator Tat, which recruits positive transcription elongation factor b to promote efficient transcriptional elongation. Its central role in HIV-1 gene expression has made TAR a prominent target for small molecule ligand discovery. Ganser et al.^[26]^ screened approximately 100,000 drug-like compounds against TAR using a Tat peptide displacement assay and subsequently validated the identified hits using dose-response and NMR-based experiments. The screen yielded seven validated TAR binders, which were subsequently combined with 170 previously reported TAR binders. To reduce structural redundancy and maximize chemical diversity, the 177 binders were clustered according to Bemis-Murcko scaffold, and the highest-affinity compound from each scaffold was retained as a representative. This procedure yielded the ‘full set’, consisting of 78 binders and 103,349 experimentally verified non-binders. The authors further noted that the hits generally had higher molecular weight and charge, more rotatable bonds and hydrogen-bond donors and acceptors, and lower logP values than non-hits. Since the differences could lead to artificial enrichment based on physicochemical properties, the authors constructed more stringent property-matched libraries. In the ‘optimized set’, experimentally verified non-binders were selected to match the physicochemical properties of the hits while maintaining topologically distinct structures, resulting in a benchmark comprising 14 binders and 637 non-hits.

The structure-based virtual screening studies, including SPRank^[10]^ and SIGNALD^[27]^, evaluated the smaller optimized set rather than the full set. We therefore compared SIREN on the full set primarily with the original ensemble screening reported by Ganser et al., while broader comparisons with other scoring methods were performed on the optimized set. On the full set, ICM docking against the experimentally informed TAR conformational ensemble (E_0,4rdc_) achieved an AUROC of 0.88 and recovered 42% of the known hits within the top 2% false-positive rate. In comparison, SIREN achieved an AUROC of 0.93 and recovered 60.3% of the known hits under the same criterion without using RNA structural information.

On the property-matched optimized set, SIREN achieved an AUROC of 0.86, the highest among the methods evaluated in **Table 2**. In comparison, the structure-based methods achieved AUROCs ranging from 0.37 to 0.79, with SIGNALD yielding the highest value (0.79) when compounds were ranked by the best score. Thus, SIREN maintained strong overall discrimination even under the more stringent property-matched benchmark without requiring an RNA tertiary structure or conformational ensemble.

**Table 2.** Virtual-screening performance on the HIV-1 TAR optimized set. Enrichment factors at the top 2%, 3%, and 5% of the ranked compound library and AUROC are reported for each scoring function under its evaluated ensemble strategy. Structure-based comparator values are taken from the SIGNALD study^[27]^; SMRTnet values were obtained in this study by applying its publicly released model to the same compound set. Five inactive molecules were removed in the SIGNALD study because rDock could not treat them.

| **Method** | **Ensemble** | **EF2%** | **EF3%** | **EF5%** | **AUROC** |
|---|---|---|---|---|---|
| SIREN | — | 3.57 | 7.14 | 4.29 | 0.86 |
| SIGNALD | Best score | 7.10 | 4.61 | 5.59 | 0.79 |
|  | Average rank | 17.75 | 13.84 | 9.79 | 0.76 |
| SPRank | Best score | 10.65 | 9.23 | 6.99 | 0.76 |
|  | Average rank | 3.55 | 4.61 | 5.59 | 0.74 |
| RmsdXNA | Best score | 0.00 | 2.31 | 1.40 | 0.63 |
|  | Average rank | 3.55 | 2.31 | 1.40 | 0.52 |
| PLANTS | Best score | 0.00 | 0.00 | 1.40 | 0.58 |
|  | Average rank | 0.00 | 0.00 | 1.40 | 0.54 |
| rDock | Best score | 0.00 | 0.00 | 0.00 | 0.40 |
|  | Average rank | 0.00 | 0.00 | 0.00 | 0.37 |
| RNAmigos2 | — | 0.00 | 2.31 | 2.78 | 0.75 |
| SMRTnet | — | 3.57 | 2.38 | 1.43 | 0.65 |

Early enrichment showed a different pattern. SIREN achieved EF2%, EF3%, and EF5% values of 3.57, 7.14, and 4.29, respectively. At the 2% cutoff, this enrichment was lower than that obtained with SIGNALD using average-rank ensemble aggregation (EF2% = 17.75) and SPRank using the best-score strategy (EF2% = 10.65). SIREN nevertheless maintained enrichment above random retrieval across all evaluated cutoffs and compared favorably with several other structure-based and structure-free approaches at the broader cutoffs. These results indicate that SIREN provides strong global discrimination on the property-matched TAR benchmark, although ensemble-based structure methods can provide greater enrichment within the very highest-ranked fraction of the library. Because the optimized set was specifically constructed to reduce simple physicochemical differences between binders and non-binders, SIREN’s performance is unlikely to be explained solely by the physicochemical descriptors incorporated into the model.

To further compare SIREN with a recently reported method that does not require RNA tertiary structure, we applied the publicly released SMRTnet model^[15]^ to the same TAR optimized set. Because SMRTnet accepts RNA inputs of 31 nucleotides, the TAR sequence was partitioned into 27 overlapping fragments, and each compound was evaluated against all fragments following the model’s default usage. SMRTnet achieved an AUROC of 0.65, compared with 0.86 for SIREN. Although the two methods showed the same EF2% (3.57), SIREN maintained higher enrichment at the broader cutoffs (EF3% = 7.14 and EF5% = 4.29) than SMRTnet (EF3% = 2.38 and EF5% = 1.43). These results further support the ability of SIREN to rank TAR-binding compounds using sequence-derived RNA representations without explicit tertiary-structure information.

To examine whether a SIREN-prioritized binder could be accommodated with an experimentally informed TAR structure, we performed a post hoc docking analysis of an experimentally validated binder, an isoquinoline derivative (IG_1_), ranked 8th by SIREN using the residual dipolar coupling (RDC)-derived HIV-1 TAR conformational ensemble originally reported by Salmon et al^[28]^. The compound binds to HIV-1 TAR RNA and thus inhibits Tat– TAR interactions^[29]^. Docking identified a favorable binding pose with an rDock score of −32.17 in one ensemble conformer (**Figure 3**). In this pose, the ligand occupied a pocket formed by C19, A20, G21, and G36 and formed several hydrogen-bond-compatible polar contacts, including a 2.67 Å contact between its positively charged ligand moiety and a phosphate oxygen of A20, without obvious steric clashes. Previous experimental studies have demonstrated that TAR-binding ligands can interact with the RNA through electrostatic contacts involving the phosphate backbone and through recognition of structural features in the bulge and adjacent stem regions^[30]^. In particular, footprinting and NMR studies have implicated residues in the lower stem, including C19, A20, and G21, as well as G36 in the upper stem, in ligand-associated perturbations^[31]^. Independent crystal structures of full-length HIV-1 TAR provide further structural support for these contacts: the bulge and flanking stem region constitute a conformationally dynamic recognition platform in which a critical Tat arginine engages the Hoogsteen edge of a bulge-proximal guanine, and divalent cations specifically coordinate phosphate oxygens of the distorted bulge backbone, paralleling the electrostatic phosphate contact observed in the SIREN-prioritized docking pose^[32]^. These structures also reveal substantial conformational heterogeneity at the bulge across independently solved copies, reinforcing the view that the docked pose represents one plausible conformer within a dynamic ensemble rather than a single, fixed binding geometry. These observations are consistent with the general interaction features of the predicted pose. Although the docking analysis does not establish the experimental binding mode, the pose demonstrates that the SIREN-prioritized ligand can be accommodated within an experimentally informed TAR conformation without obvious steric incompatibility.

**Figure 3.**
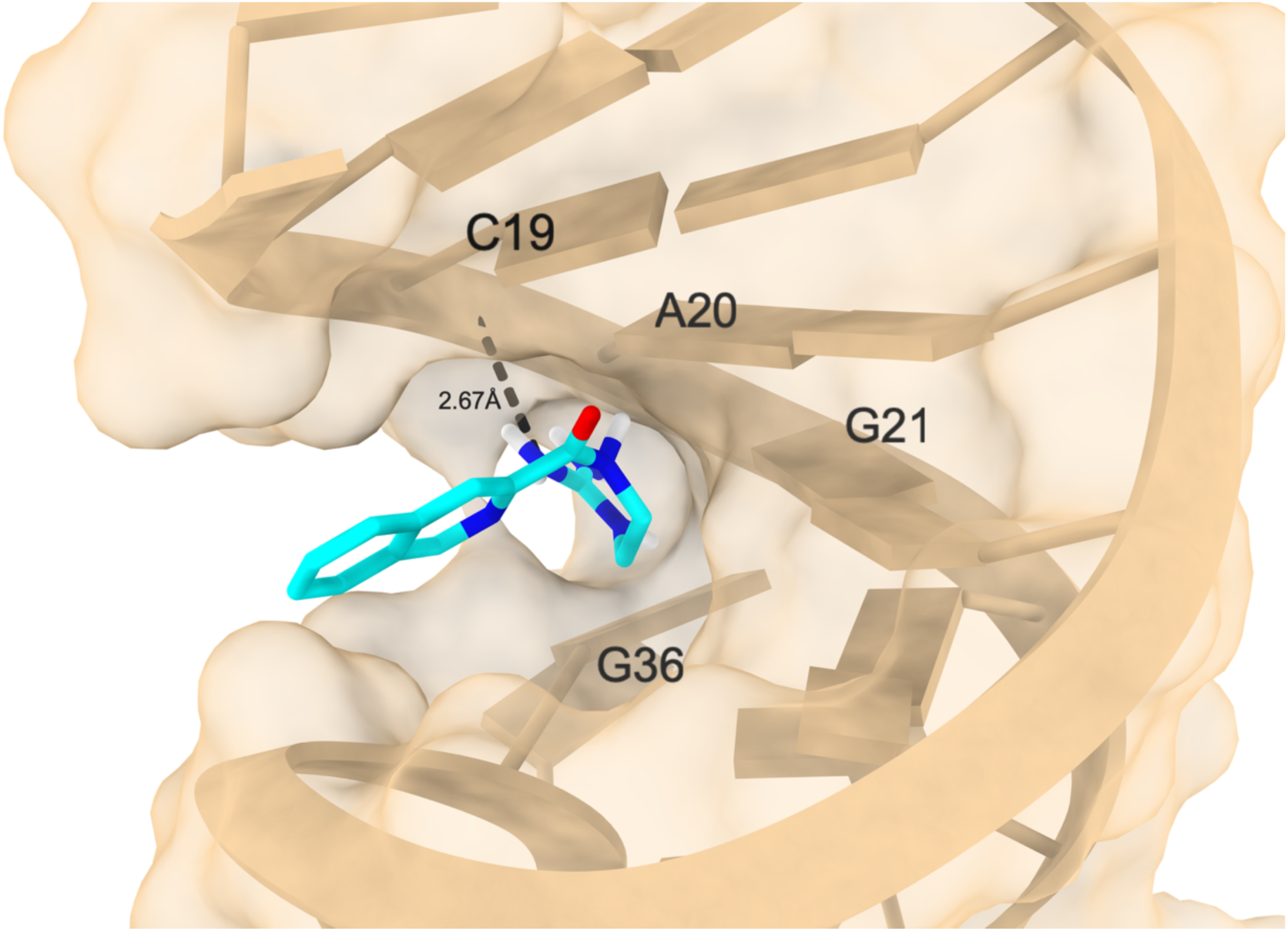
Predicted binding pose of a SIREN-prioritized HIV-1 TAR binder using ensemble docking. The dashed black line indicates an interaction between the positively charged ligand moiety and the A20 phosphate oxygen.

Using precomputed RNA and ligand representations, SIREN scored the TAR benchmark compounds in 54.5 s on a single 4070 Ti Super, corresponding to approximately 1,898 compounds per second. For context, SPRank reported approximately 17.5 h for docking and scoring the 651-compound optimized library against the 20-conformer TAR ensemble.

### 2.5 Experimental validation of SIREN-prioritized compounds against cMyc and XIAP IRES-derived RNA hairpins

To assess the practical utility of SIREN for prospective screening, the model was applied to two independent cancer-associated RNA targets, cMyc and XIAP mRNAs. The two targets contain structured internal ribosome entry sites (IRES) that contribute to cap-independent translation and therefore represent functionally relevant RNA elements for ligand discovery. Previous studies have characterized the secondary structures of the cMyc and XIAP IRES regions, revealing structured motifs that may serve as potential small-molecule binding sites ^[33,34]^. Despite this experimental support for their secondary structures, high-resolution tertiary structures of these RNAs remain unavailable. Thus, cMyc and XIAP provide suitable prospective targets for evaluating SIREN in a setting where experimentally resolved RNA structures are not available. The RNA sequences used for screening are provided in **Table S3**.

For each RNA target, SIREN screened 2,333 compounds from the ChemBank of KRICT. After ranking the compounds according to the SIREN score, the top 10% of the library (233 compounds) per target were selected and further experimentally validated using two complementary fluorescence-based assays for each target. The cMyc IRES-derived hairpin was evaluated using a Cy5-labeled RNA quenching assay and a TO-PRO-3 displacement assay. The XIAP IRES-derived hairpin was evaluated using a 2-aminopurine (2-AP) fluorescence assay, with 2-AP incorporated at the adenine bulge, and a TO-PRO-3 displacement assay. These assays provided distinct readouts of compound-induced changes in the local RNA environment or fluorophore fluorescence. Compounds exhibiting a fluorescence response in at least one of the two assays were designated as putative RNA binders. Of the 233 compounds experimentally tested for each target, 13 and 14 were identified as binders for cMyc and XIAP IRES-derived hairpins, respectively. Binding affinities of these hits were subsequently quantified by measuring their dissociation constants (K_D_). Notably, four compounds targeting cMyc and three compounds targeting XIAP exhibited sub-10 μM dissociation constants (K_D_ ≤ 10 μM), with an additional XIAP-binding compound reaching K_D_ ≤ 15 μM. **Figure 4** shows their two-dimensional structures and concentration-response curves. These results demonstrated that SIREN prioritized experimentally confirmed, micromolar-affinity binders for both RNA targets from the top-ranked screening subset. The detailed assay results of the selected 233 compounds for each target RNA are given in the Supporting Information.

**Figure 4.**
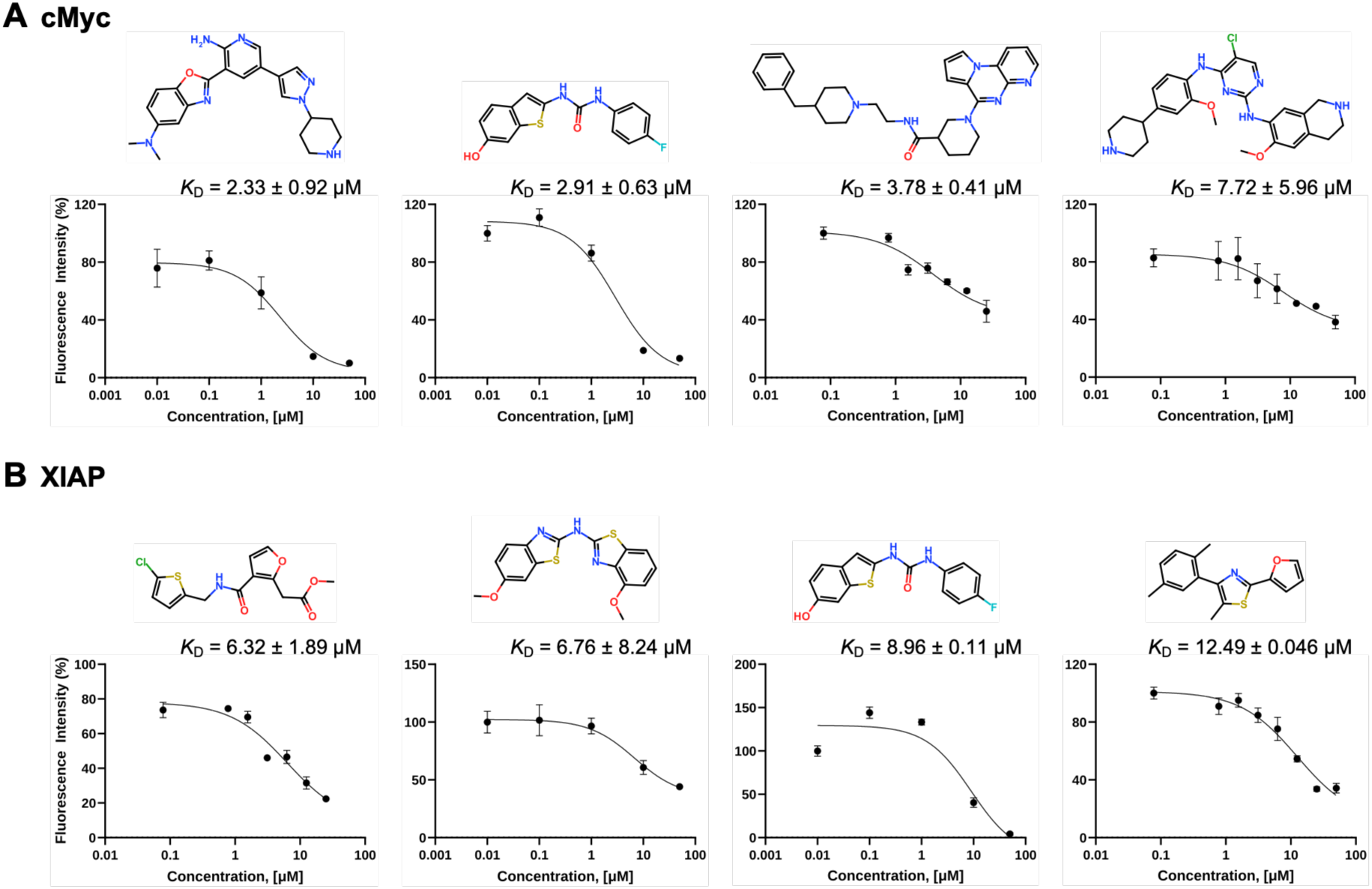
Chemical structures and concentration-response curves of experimentally validated SIREN-prioritized binders for (A) cMyc and (B) XIAP RNAs. The K_D_ values, together with their reported uncertainties, are reported in the upper right corner of each plot.

## 3. Discussion

SIREN demonstrates that primary RNA sequence can provide sufficient information for RNA-targeted virtual screening without requiring experimentally resolved tertiary structures or predefined binding pockets. A systematic evaluation of RNA and ligand representations identified RiNALMo combined with ChemBERTa and Mordred descriptors as the best-performing configuration by validation AUPRC. The observed tendency for descriptor-containing ligand representations to outperform molecular language-model embeddings alone is consistent with the complementarity of the two feature types in capturing ligand chemistry. RiNALMo’s consistently strong performance across ligand representations suggests that pretrained RNA sequence embeddings contribute information relevant to ligand discrimination. In a broader sense, SIREN was evaluated beyond conventional pairwise classification through held-out RNA screening, target-swapping analysis, and an independent external benchmark, and prospective experimental validation, providing multiple levels of evidence for its utility as a screening framework.

A central question for sequence-based RNA-ligand models is whether their predictions genuinely depend on RNA identity or instead reflect general ligand-binding propensity. On four ROBIN targets excluded from model development, SIREN achieved higher mean AUROC and substantially greater early enrichment compared to the reported RNAmigos2, RNAmigos++, and rDock benchmarks. The gains were particularly pronounced among the top 1%, 2%, and 5% of ranked libraries. However, the cross-target derangement analysis provides a more direct test of target dependence. The reassignment of prediction scores among RNA targets significantly reduced both pooled AUROC and EF1%, indicating that the model uses RNA-specific information. The magnitude of this effect was moderate, suggesting that SIREN combines target-dependent recognition with a broader ligand-propensity component. This distinction is important because recent analyses of sequence-based RNA– ligand models have shown that apparently strong predictive performance can arise in part from target-independent ligand signals^[35]^. The ROBIN results should also be interpreted within the context of SMM measurements rather than as direct estimates of solution-phase binding affinity.

The HIV-1 TAR benchmark further illustrates both the strength and limitations of a structure-free screening strategy. On the full set, SIREN attained an AUROC of 0.93. It recovered 60.3% of known hits within the top 2% false-positive rate, compared with an AUROC of 0.88 and 42% recovery at the same cutoff for ensemble-based docking against an experimentally informed E_0,4rdc_ TAR ensemble. This comparison suggests that sequence-derived representations can support large-scale ligand ranking without needing an experimentally determined conformational ensemble. However, the full set contains known physicochemical differences between binders and non-binders, and performance on this benchmark may therefore partly reflect ligand-level property signals.

The property-matched TAR optimized set provides a more stringent assessment because it was designed to reduce simple physicochemical differences between binders and non-binders. On this benchmark, SIREN achieved the highest overall discrimination among the evaluated methods (AUROC = 0.86), exceeding both ensemble-based structure methods and the structure-free RNAmigos2 and SMRTnet models. This result is particularly relevant because SIREN explicitly incorporates molecular descriptors, yet its performance remained strong when simple physicochemical differences between the two classes were minimized. Together with the cross-target analysis, this finding suggests that SIREN captures screening information beyond a general preference for compounds with particular physicochemical properties.

The enrichment results, however, reveal a complementary strength of structure-based screening. SIREN showed moderate enrichment at the top 2% of the optimized library (EF2% = 3.57), whereas the strongest ensemble-based strategies achieved substantially greater enrichment at this cutoff. SIREN nevertheless retained enrichment at the 3% and 5% cutoffs and outperformed several other structure-based and structure-free approaches. Thus, sequence-derived representations appear effective for global library prioritization, whereas explicit structural information can provide an advantage in resolving the very highest-ranked candidates. These observations support a hierarchical screening strategy in which SIREN rapidly reduces large chemical libraries before more computationally intensive docking or structural rescoring is applied to the prioritized subset.

The computational efficiency of SIREN is a particularly advantageous feature in a hierarchical screening workflow. Once RNA and ligand representations were precomputed, SIREN scored more than 100,000 TAR compounds in less than one minute on a single GPU. In contrast, ensemble docking requires evaluating each ligand against multiple RNA conformations. This contrast highlights the superior scaling of representation-based screening compared with explicit ensemble docking.

Virtual screening against IRES-derived RNA hairpins from cMyc and XIAP mRNAs provides evidence that SIREN can prioritize experimentally measurable binders for previously unseen human RNA targets. Both targets contain functionally relevant IRES elements with experimentally supported secondary structures, but high-resolution tertiary structures remain unsolved. This renders them representative use cases for a sequence-based screening method. SIREN ranked 2,333 compounds for each RNA, and the top 10% of these compounds were subjected to experimental testing. We identified multiple binders for both RNAs, including four cMyc-binding compounds and three XIAP-binding compounds with K_D_ ≤ 10 μM, plus one additional XIAP-binding compound with K_D_ ≤ 15 μM. The recovery of the micromolar binders for two disease-associated human RNAs extends the evaluation beyond retrospective riboswitch and viral benchmarks and supports the practical utility of the model in settings where experimentally determined RNA structures are unavailable.

SIREN has several limitations despite its strong performance in screening RNA-targeting ligands. SIREN does not explicitly model RNA structure, conformational dynamics, or induced fit. Its output should be interpreted as a ranking score rather than a calibrated affinity estimate. In addition, the moderate effect observed in the cross-target derangement analysis suggests that predictions retain a target-independent ligand component and may reflect biases in the training data. Future work should therefore focus on broader prospective validation and integration with structural information or structure-based rescoring. Overall, SIREN provides a scalable sequence-based framework for RNA-targeted compound prioritization that complements structure-based screening.

## 4. Methods and Experiments

### 4.1. Architecture of SIREN

SIREN is a neural network to predict RNA–ligand interactions from pretrained RNA and ligand representations without requiring RNA secondary-or tertiary-structure information. The final configuration utilizes RiNALMo^[18]^ to encode RNA sequences and combines ChemBERTa^[21]^ embeddings with Mordred molecular descriptors to represent ligands. Because RiNALMo supports input sequences of up to 1,022 nucleotides, SIREN is currently limited to RNA sequences within this length.

Contextualized nucleotide-level RNA embeddings generated by RiNALMo were projected into a 256-dimensional latent space using a linear layer followed by layer normalization, Gaussian error linear unit (GELU) activation, and dropout. A learnable attention module assigned a scalar weight to each nucleotide embedding, and masked softmax normalization was applied to exclude padded positions. The weighted nucleotide representations were subsequently aggregated to obtain a fixed-dimensional RNA vector *r*.

Ligands were represented as fixed-length vectors. For multimodal ligand representations, the ChemBERTa embedding and Mordred descriptor vector were concatenated and projected into the same 256-dimensional latent space to obtain a ligand vector *l*. Because each ligand was represented by a single fixed-length feature vector, the pooling operation reduced to the projected ligand representation itself.

RNA–ligand compatibility was modeled by separately projecting the RNA and ligand vectors and combining the resulting representations through element-wise multiplication (**Equation 1**).

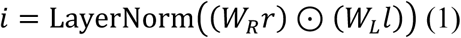

The final pair representation was constructed by concatenating the RNA vector, ligand vector, and their element-wise absolute difference, and the multiplicative interaction vector (**Equation 2**).

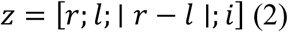

The resulting 1,024-dimensional pair representation was projected into a 512-dimensional hidden space using a linear layer followed by layer normalization, GELU activation, and dropout. The hidden representation was then processed by two residual multilayer perceptron blocks. Each block consisted of pre-layer normalization and a two-layer feed-forward network that expanded the representation from 512 to 1,024 dimensions and projected it back to 512 dimensions, followed by residual addition. A dropout rate of 0.15 was applied throughout the interaction network. Finally, the refined representation was mapped to a scalar interaction logit through a linear output layer. The logits were transformed using a sigmoid function, and the resulting scores were used to rank candidate ligands for each RNA target.

Separate auxiliary prediction heads were applied to the pooled RNA and ligand representations to estimate RNA-level and ligand-level interaction propensities, respectively. Each head consisted of layer normalization followed by a two-layer multilayer perceptron. The auxiliary predictions were used only as an additional training objective and did not contribute to the final RNA–ligand interaction score.

### 4.2. RNA Representation using Language Models

Pretrained RNA language models were used to generate sequence representations for SIREN. Recent studies^[14,36]^ have shown that residue-level embeddings derived from RNA language models can support nucleotide-level small-molecule binding site prediction directly from primary RNA sequence without requiring explicit structural information.

Three pretrained RNA language models were evaluated in this study: ERNIE-RNA^[19]^, RiNALMo^[18]^, and RNA-FM^[20]^. ERNIE-RNA is a BERT-style architecture that incorporates structural information through an attention-bias mechanism. RiNALMo is a 650-million-parameter Transformer encoder pretrained on approximately 36 million noncoding RNA sequences using masked language modeling. RNA-FM is a 12-layer Transformer encoder pretrained on approximately 23 million noncoding RNA sequences using a masked language-modeling objective. All three encoders operate directly on primary RNA sequence without requiring multiple sequence alignments or explicit tertiary structure information. For each model, contextualized nucleotide-level embeddings were extracted and used as the RNA input representations for SIREN.

### 4.3. Small Molecule Representations using Language Models and Molecular Descriptors

Small molecules were represented using pretrained molecular language models and physicochemical descriptors. Two Transformer-based molecular language models operating on SMILES strings, ChemBERTa-77M-MTR^[21]^ and MoLFormer-XL-both-10pct^[22]^, were evaluated. ChemBERTa is based on the RoBERTa architecture and consists of three Transformer layers with 12 attention heads. The model was pretrained on approximately 77 million canonical SMILES strings derived from PubChem. MoLFormer is a Transformer encoder incorporating linear attention and rotary positional embeddings. The MoLFormer-XL configuration consists of 12 Transformer layers with 12 attention heads and a hidden dimension of 768 and was pretrained using masked language modeling on 10% of subsets of both PubChem and ZINC databases. For each compound, representations from the final hidden layer were mean-pooled across all non-padding tokens, including special tokens, to obtain a fixed-length molecular embedding, which was subsequently used as a ligand feature vector in SIREN.

In parallel, molecular descriptors were calculated using Mordred^[24]^ and RDKit^[23]^. Before descriptor calculation, multicomponent SMILES strings were standardized by retaining the largest carbon-containing fragment, thereby removing smaller salts and counterions. Protonation states were then adjusted to approximate physiological pH (7.4) using Dimorphite-DL^[37]^. When protonation-state generation failed, the standardized input SMILES was retained.

For descriptors requiring three-dimensional information, a conformer was generated from each processed SMILES using the ETKDGv3 algorithm with explicit hydrogens and chirality constraints. Conformers were optimized using MMFF94s when suitable force-field parameters were available and UFF otherwise. The resulting conformer was used to calculate Mordred descriptors and the complete sets of RDKit 2D and 3D descriptors. Descriptor values that could not be calculated or were non-finite were treated as missing. Features with more than 40% missing values in the training set were removed, and remaining missing values were imputed using the corresponding training-set median. Retained descriptor features were standardized using training-set means and standard deviations and clipped to [−10, 10] after standardization.

### 4.4 Training Objective and Optimization

SIREN was trained using two complementary batch-sampling strategies designed to learn both within-target ligand ranking and ligand selectivity across RNA targets. RNA-centered batches were constructed by sampling eight RNA targets per batch. For each RNA, four positive ligands were sampled together with four negative ligands per positive ligand, yielding four positive and sixteen negative RNA–ligand pairs per target. Ligand-centered batches were generated separately by sampling eight ligands per batch, with one RNA target sampled from each ligand’s positive target set and four negative RNA targets sampled for each ligand.

For RNA-centered batches, the training objective combined binary interaction classification with a within-RNA ranking loss. The ranking loss encouraged experimentally positive ligands to receive higher interaction scores than the negative ligands for the same RNA target. Hard-negative mining was applied by retaining up to eight of the highest-scoring negative ligands for each RNA. For ligand-centered batches, an analogous selectivity loss encouraged each ligand to receive a higher score for its experimentally positive RNA target than for negative RNA targets. Up to four highest-scoring negative RNA partners were retained for each ligand. Both ranking objectives were implemented using a margin-based hinge loss with a margin of 0.2.

In addition, RNA-and ligand-level auxiliary heads were trained to predict empirical interaction propensities derived from the training data. The RNA-level propensity target was defined as the fraction of positive interactions among all training pairs associated with a given RNA, whereas the ligand-level propensity target was defined analogously across RNA partners for each ligand. Sigmoid-transformed auxiliary predictions were optimized against these targets using smooth L1 loss. These auxiliary losses were used only during training and propagated gradients through the corresponding trainable projection and pooling modules.

At each optimization step, the RNA-centered batch contributed the binary cross-entropy loss, within-RNA ranking loss, and RNA-and ligand-propensity losses, whereas the ligand-centered batch contributed the ligand-selectivity loss. The total training objective was defined as **Equation 3**.

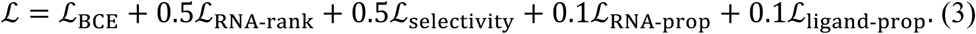

Five hundred RNA-centered and 500 ligand-centered batches were generated per epoch, with one batch of each type jointly contributing to each optimization step. Model parameters were optimized using AdamW with an initial learning rate of 1×10^-3^ and a weight decay of 1×10^-4^. Training was performed for up to 50 epochs using mixed-precision computation, with gradient norms clipped to 5.0. The learning rate was reduced by a factor of 0.5 after two consecutive epochs without improvement in validation AUPRC, with a minimum learning rate of 1×10^-6^. Early stopping was applied after eight consecutive epochs without improvement, and the checkpoint with the highest validation AUPRC was retained for subsequent evaluation.

### 4.5 RNA-Small-Molecule Datasets for Model Development and Benchmarking

#### 4.5.1. ROBIN Dataset and Data Partitioning

The ROBIN dataset was obtained from a previously published SMM study^[17]^. In the SMM assay, chemically diverse small molecules were immobilized on microarray surfaces and incubated with fluorescently labeled biomolecular targets, with binding assessed from fluorescence signal intensity. The dataset contains measurements for nine DNA and 27 RNA targets, each screened against a library of 24,572 small molecules. Only the 27 RNA targets were utilized in this study. RNA–compound pairs were assigned binary interaction labels by applying predefined signal thresholds to the reported SMM measurements; quantitative binding affinities were not available.

Four RNA targets—PreQ1, SAM-II, TPP, and ZTP—previously evaluated in RNAmigos2^[12]^ were reserved as a fixed held-out test set and were excluded from model training and selection. The remaining RNA targets were partitioned into training and validation sets on the basis of sequence similarity to reduce sequence similarity between the two subsets. Pairwise sequence similarity was calculated from the normalized Levenshtein distance (**Equation 4**), where the Levenshtein distance corresponds to the minimum number of substitutions, insertions, and deletions (*d_lev_*)required to transform one sequence into another.

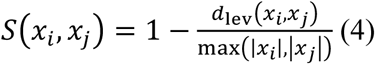

RNA pairs satisfying the similarity criterion, *S(x_i_, x_j_)* ≥ 0.8, were connected in a graph, and each connected component was treated as an indivisible sequence group. Consequently, directly or transitively related RNA sequences were assigned to the same partition.

The resulting sequence groups were divided into training and validation sets, with approximately 20% of the development RNA targets assigned to validation. To obtain a representative split while preserving sequence-group separation, 10,000 randomized cluster-level candidate splits were generated. The final split was selected to approximate the target validation fraction while minimizing differences in positive-interaction prevalence between the training and validation sets. The four held-out test RNAs were not considered during this procedure and were used for final evaluation.

#### 4.5.2. HIV-1 TAR Benchmark Dataset

An independent HIV-1 TAR benchmark dataset reported by Ganser et al.^[26]^ was used to evaluate SIREN. The original study screened approximately 100,000 drug-like compounds against HIV-1 TAR and subsequently combined experimentally validated screening hits with previously reported TAR binders. The resulting 177 candidate binders were clustered according to Bemis-Murcko frameworks, and the highest-affinity compound from each scaffold was retained to reduce structural redundancy. This procedure yielded 78 structurally distinct binders. The full set comprised these 78 binders together with 103,349 experimentally verified non-binders, resulting in a total of 103,427 compounds.

Because binders and non-binders in the full set differed in several physicochemical properties, Ganser et al. additionally constructed a more stringent property-matched benchmark. In this optimized set, experimentally verified non-binders were selected to approximate the physicochemical properties of the binders while remaining topologically distinct. The resulting dataset contained 14 binders and 637 non-binders, for a total of 651 compounds.

SIREN was evaluated on both benchmark sets: the full set was used for comparison with the original ensemble-based virtual screening study, whereas the optimized set was used for comparisons with structure-based scoring methods. Neither the full nor the optimized TAR dataset was used during SIREN training or model selection.

For comparison, performance metrics for the structure-based methods were taken directly from Tan et al.^[27]^. For SMRTnet^[15]^, we installed the stable release and ran it on Google Colab. The secondary structure, represented as dot-bracket notation, was predicted by the RNAcomp2D web server ^[38]^ and supplied as the input to the model.

### 4.6 Evaluation Metrics and Statistical Analysis

Model performance was evaluated using AUPRC, AUROC, and EFs. AUPRC was used as the primary metric because of the strong class imbalance in RNA-ligand interaction data, while AUROC was reported as a complementary measure of overall discrimination. Early retrieval performance was assessed using EF at the top 1%, 2%, and 5% of each ranked compound library. EFs at the top *x*% is calculated as in **Equation 5**.

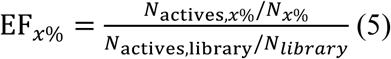

*N_library_* and *N_active_* denote the total number of compounds and active molecules in the screened library, respectively, and *N_x%_* and *N_actives,x%_* denote the corresponding numbers within the top *x*% of ranked library. An EF of 1 corresponds to random retrieval, whereas values greater than 1 indicate enrichment of binders among the top-ranked compounds.

RNA-target specificity was assessed using the 14,302 ligands shared across the four held-out ROBIN targets. The correctly matched RNA–score assignment was compared with all nine complete cross-target derangements, in which no RNA retained scores generated for itself. AUROC was calculated using within-target percentile ranks to account for differences in score distributions across targets, whereas EF1% was calculated from raw prediction scores. The specificity effect was defined as the difference between the metric obtained for the correct assignment and the mean across the nine derangements.

Statistical uncertainty was estimated using 2,000 ligand-level bootstrap replicates, with each sampled ligand retaining its observations across all four RNA targets. Ninety-five percent confidence intervals were obtained from the 2.5th and 97.5th percentiles of the bootstrap distribution, and one-sided *P*-values were calculated as the proportion of bootstrap differences less than or equal to zero using an add-one correction.

### 4.7 Ensemble Docking of a SIREN-Prioritized TAR Binder

To assess whether a SIREN-prioritized, experimentally validated TAR binder could be accommodated within experimentally informed TAR conformations, ensemble docking was performed against the 20-conformer RDC-derived HIV-1 TAR ensemble reported by Salmon et al.^[28]^. Docking was conducted using rDock v25.07-alpha^[25]^. Each RNA conformer was converted from PDB to MOL2 format using OpenBabel v3.2.1^[39]^, followed by standardization of RNA residue and phosphate atom names. The ligand was prepared from SMILES using RDKit^[23]^. Explicit hydrogen atoms were added, and a three-dimensional conformer was generated using ETKDGv3. Geometry optimization was performed using MMFF94 when suitable parameters were available and UFF otherwise, with a maximum of 1,000 iterations.

For each conformer, the docking box was defined using the RbtSphereSiteMapper, with the search sphere centered at the geometric center of the RNA and extending 8 Å beyond the maximum atomic distance from the center. Cavity mapping was performed using SMALL_SPHERE = 1.5 Å, LARGE_SPHERE = 4.0 Å, MAX_CAVITIES = 99, and MIN_VOLUME = 100 Å³ with the RbtCavityGridSF scoring function. The ligand was independently docked to each of the 20 conformers using 500 docking runs per conformer. The lowest-scoring pose across the ensemble was selected for structural visualization and contact analysis. RNA–ligand contacts were identified using a 4.0-Å heavy-atom distance cutoff, and hydrogen-bond-compatible donor–acceptor contacts were defined using a distance cutoff of 3.5 Å. Contacts between positively charged ligand groups and RNA phosphate atoms were examined separately. Steric compatibility was assessed from the intermolecular repulsive component of the rDock score.

### 4.8 Experimental validation of cMyc- and XIAP-prioritized compounds

The 233 SIREN-prioritized compounds for each RNA, cMyc and XIAP, were evaluated using fluorescence-based assays. Cy5 fluorescence quenching and TO-PRO-3 displacement assays were employed for cMyc RNA, whereas 2-AP fluorescence and TO-PRO-3 displacement assays were used for XIAP RNA.

All RNAs used in this study were folded in folding buffer (20 mM HEPES, pH 7.5, and 50 mM NaCl) by heating at 100 ℃ for 1 min, followed by cooling at room temperature for 10 min. The RNAs were subsequently heated at 60 ℃ for 5 min and cooled at room temperature for 20 min.

For Cy5 fluorescence quenching assay, folded 5’-Cy5-cMyc RNA was diluted to a final concentration of 10 nM and incubated with increasing concentrations of the compounds.

In the TO-PRO-3 displacement assay, folded cMyc and XIAP RNAs were diluted to a final concentration of 10 nM. TO-PRO-3 iodide (Thermo Fisher Scientific, Cat. No. T3605) was added to a final concentration of 50 nM, followed by incubation at room temperature for 5 min. Compounds were subsequently added at various concentrations. The 2-AP fluorescence assay used 2-AP-labeled XIAP RNA prepared using the same folding procedure and diluted to a final concentration of 100 nM. The folded RNA was then incubated with increasing concentrations of the compounds.

For all assays, reaction mixtures (20 µL per well) were transferred to black 384-well plates (Greiner Bio-One, Cat. No. 781209) and incubated at room temperature for 30 min. Fluorescence intensities were measured using an Infinite M Plex microplate reader (Tecan) at excitation/emission wavelengths of 650/700 nm for the Cy5 fluorescence quenching assay, 600/650 nm for the TO-PRO-3 displacement assay, and 310/365 nm for the 2-AP fluorescence assay. Concentration–response curves were fitted by nonlinear regression using the [Inhibitor] vs. response (three-parameter) model in GraphPad Prism 10.

The following RNA oligonucleotides were purchased from SFC, except for the 2-AP-labeled XIAP RNA, which was purchased from Integrated DNA Technologies (IDT). 2-AP indicates the site of 2-aminopurine incorporation. The nucleic acid sequences used for the assay are listed in Table S3.

## Acknowledgements

The chemical library used in this study to discover cMyc-and XIAP-binding molecules was kindly provided by the Korea Chemical Bank (www.chembank.org) of the Korea Research Institute of Chemical Technology.

## Data Availability Statement

The benchmark datasets and data access instructions are available at https://github.com/kucm-lsbi/SIREN/tree/main/dataset. SIREN code is also available at GitHub repository, https://github.com/kucm-lsbi/SIREN.

Received: ((will be filled in by the editorial staff)) Revised: ((will be filled in by the editorial staff)) Published online: ((will be filled in by the editorial staff))

## Funding

This work is supported by Institute of Information & Communications Technology Planning & Evaluation (IITP)—ICT Challenge and Advanced Network of HRD (ICAN) grant funded by the Korea government (Ministry of Science and ICT) (IITP-2026-RS-2024-00438263); the Bio & Medical Technology Development Program of the National Research Foundation (NRF) funded by the Korea government (Ministry of Science and ICT) (RS-2025-02217289); Korea Institute for Advancement of Technology (KIAT) grant funded by the Korea Government (MOTIE) (RS-2025-02214034, HRD Program for Industrial Innovation); the Global–Learning & Academic Research Institution for Master’s, PhD Students, and Postdocs (LAMP) Program of the NRF funded by the Ministry of Education (RS-2023-00301938)

## Supporting Information

Supporting Information is available from the Wiley Online Library or from the author.

The table of contents entry should be 50–60 words long and should be written in the present tense. The text should be different from the abstract text.

## SIREN: Predicting RNA-Small Molecule Interaction using Language Models without Requiring Structure

SIREN is a sequence-based virtual screening framework that ranks small molecules for RNA targets without requiring tertiary structures. It combines a pretrained RNA language model with molecular language model embeddings and descriptors. SIREN outperforms structure-based methods on held-out riboswitches, generalizes to HIV-1 TAR, and prioritizes experimentally validated binders for cMyc and XIAP IRES hairpins.

## ToC figure

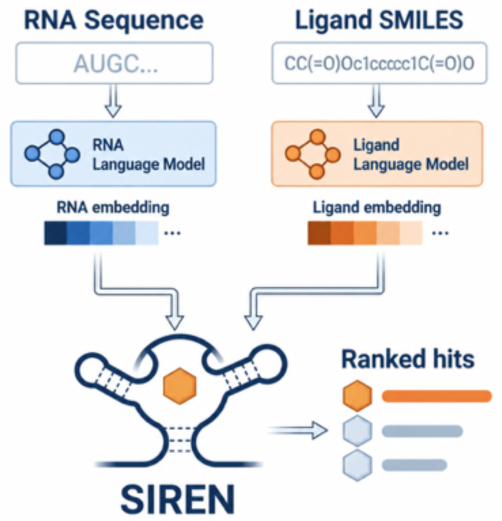

## Supporting Information

**Table S1.** Validation performance across 24 configurations. AUPRC and AUROC are reported for all combinations of RNA language models and ligand representation configurations.

| **RNA Language Model** | **Ligand Representation** | **AUPRC** | **AUROC** |
|---|---|---|---|
| ERNIE-RNA | ChemBERTa | 0.065 | 0.749 |
|  | ChemBERTa+Mordred | 0.101 | 0.769 |
|  | ChemBERTa+RDKit | 0.095 | 0.796 |
|  | MolFormer | 0.080 | 0.747 |
|  | MolFormer+Mordred | 0.092 | 0.776 |
|  | MolFormer+RDKit | 0.080 | 0.768 |
|  | Mordred | 0.093 | 0.775 |
|  | RDKit | 0.080 | 0.755 |
| RNA-FM | ChemBERTa | 0.066 | 0.722 |
|  | ChemBERTa+Mordred | 0.085 | 0.763 |
|  | ChemBERTa+RDKit | 0.072 | 0.745 |
|  | MolFormer | 0.089 | 0.739 |
|  | MolFormer+Mordred | 0.093 | 0.786 |
|  | MolFormer+RDKit | 0.090 | 0.763 |
|  | Mordred | 0.086 | 0.753 |
|  | RDKit | 0.074 | 0.743 |
| RiNALMo | ChemBERTa | 0.082 | 0.758 |
|  | ChemBERTa+Mordred | 0.111 | 0.772 |
|  | ChemBERTa+RDKit | 0.092 | 0.805 |
|  | MolFormer | 0.095 | 0.749 |
|  | MolFormer+Mordred | 0.100 | 0.765 |
|  | MolFormer+RDKit | 0.097 | 0.802 |
|  | Mordred | 0.097 | 0.742 |
|  | RDKit | 0.090 | 0.765 |

**Table S2.** Target-specific virtual-screening performance on the held-out ROBIN benchmark.

| Method | SIREN | RNAmigos2 | RNAmigos++ | rDock |
|---|---|---|---|---|
| PreQ1 |  |  |  |  |
| EF1% | 9.14 | 1.80 | 3.59 | 4.79 |
| EF2% | 8.22 | 0.90 | 3.29 | 3.89 |
| EF5% | 5.24 | 1.68 | 2.40 | 3.47 |
| AUROC | 0.67 | 0.62 | 0.65 | 0.61 |
| SAM-II |  |  |  |  |
| EF1% | 12.80 | 5.09 | 4.58 | 5.60 |
| EF2% | 9.09 | 3.30 | 5.08 | 4.83 |
| EF5% | 5.88 | 2.64 | 3.45 | 3.45 |
| AUROC | 0.71 | 0.66 | 0.68 | 0.66 |
| TPP |  |  |  |  |
| EF1% | 15.18 | 3.68 | 5.52 | 6.14 |
| EF2% | 12.33 | 1.84 | 4.30 | 3.68 |
| EF5% | 8.23 | 2.33 | 2.58 | 2.95 |
| AUROC | 0.77 | 0.62 | 0.65 | 0.62 |
| ZTP |  |  |  |  |
| EF1% | 16.02 | 1.17 | 0.00 | 1.76 |
| EF2% | 11.00 | 0.88 | 1.75 | 4.09 |
| EF5% | 6.78 | 2.22 | 2.11 | 2.93 |
| AUROC | 0.71 | 0.61 | 0.61 | 0.63 |

**Table S3.**
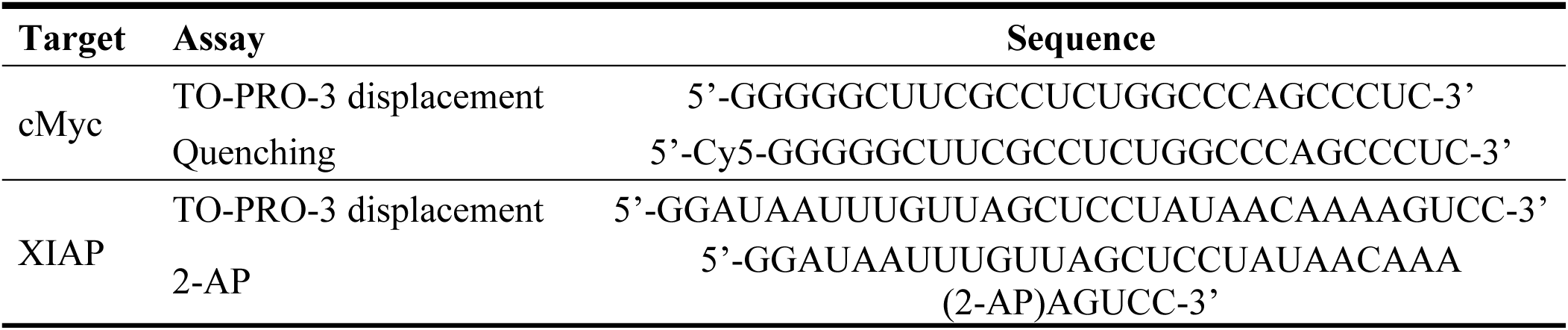
RNA sequences utilized for binding assay of SIREN-prioritized molecules.

## Notes

### Competing Interest Statement

The authors have declared no competing interest.

